# A convolutional neural network for predicting transcriptional regulators of genes in Arabidopsis transcriptome data reveals classification based on positive regulatory interactions

**DOI:** 10.1101/618926

**Authors:** Dan MacLean

**Affiliations:** The Sainsbury Laboratory, University of East Anglia, Norwich Research Park, Norwich, UK, NR4 7UH

## Abstract

Gene Regulatory networks that control gene expression are widely studied yet the interactions that make them up are difficult to predict from high throughput data. Deep Learning methods such as convolutional neural networks can perform surprisingly good classifications on a variety of data types and the matrix-like gene expression profiles would seem to be ideal input data for deep learning approaches. In this short study I compiled training sets of expression data using the Arabidopsis AtGenExpress global stress expression data set and known transcription factor-target interactions from the Arabidopsis PLACE database. I built and optimised convolutional neural networks with a best model providing 95 % accuracy of classification on a held-out validation set. Investigation of the activations within this model revealed that classification was based on positive correlation of expression profiles in short sections. This result shows that a convolutional neural network can be used to make classifications and reveal the basis of those calssifications for gene expression data sets, indicating that a convolutional neural network is a useful and interpretable tool for exploratory classification of biological data. The final model is available for download and as a web application.

## Introduction

Gene regulatory networks are molecular interaction networks that control the expression of genes. These networks play essential roles in all aspects of cellular activity, acting as integrators of cell signalling pathways and acting as one layer of control of the abundance of necessary proteins in the cell. The transcriptional component of these networks comprises the core transcriptional machinery and numerous condition specific protein transcription factors that bind promoters of target genes and affect, positively or negatively, the rate of transcription of the gene. Further control of the effect of the gene and its protein can be mitigated by modification of the translation rate, protein processing and other biochemical states downstream.

Such networks have therefore been a topic of much study, the wide range of permutations of transcription factor target gene interactions means experimentally cataloguing them is expensive and time consuming, though high-throughput methods do exist and are among the most reliable data sources. Nonetheless the difficulty of these methods have inspired efforts to infer networks from more tractable and easily performed experiments. Some notable areas have been the prediction of transcription factor and target gene relationships *de novo* from models created by inference from data such as DNA sequence of target genes, binding experiments such as ChIPSeq and transcript abundance data from microarrays or RNAseq (Wille et al., 2004, Friedman (2004), Butte et al. (2000), Liang, Fuhrman and Somogyi (1998)). These tools integrate various information including known binding site information, expression levels and co-expression profiles to predict regulatory interactions and assemble entire networks.

The AtGenExpress global stress expression data set (Kilian et al., 2007) is a compendium of transcript expression studies carried out on the model plant *Arabidopsis thaliana* during various abiotic stress challenges including cold, drought, genotoxic, osmotic, oxidative, salt, UV-B and wounding. The data set was generated with the Affymetrix ATH1 gene chip (Redman et al., 2004) which contains probesets representing approximately 23750 genes. This chip has been widely used with over 14000 samples using it submitted to the GEO expression omnibus (Soboleva et al., 2012). There are numerous databases of experimentally demonstrated transcription factor and targets in Arabidopsis, such as AGRIS (Palaniswamy et al., 2006), PLACE and PlantCARE. The AtRegNet database contains around 4000 confirmed direct regulatory interactions.

Deep learning models based on neural networks have seen surprisingly good results in varied classification problems in recent years. Convolutional Neural Networks (CNNs) are a subclass of neural network that have been applied in image classification and facial recognition (Krizhevsky, Sutskever and Hinton, 2012, Lawrence et al. (1997)), drug discovery (Wallach, Dzamba and Heifets, 2015), time series data (Pyrkov et al., 2018) and natural language processing (Collobert and Weston, 2008). CNNs operate by composing local features into a larger hierarchical model. They achieve this by convolution of input data, essentially restriction of the input into smaller filters that are transformed into an output feature map (Lawrence et al., 1997). The application of these convolution layers downsamples the data while retaining pattern information and creates filter hierarchies by looking at proportionally larger sections of the input. Training of these models requires a classified set of data with positive and negative examples such that the model can learn to discriminate. The expression profiles in the AtGenExpress data and the known interactions in AtRegNet provide data from which such classified training sets can be made. As a proof-of-technology experiment I constructed a CNN from these data. The resulting model has strong predictive power and I document the model development and an associated web-tool below. The model can be used to predict *Arabidopsis* TF/gene relationships in abiotic stress and may be useful for researchers investigating gene function in abiotic stress.

## Methods

### Preparation of AtGenExpress abiotic stress microarray data

Affymetrix .`cel` files were downloaded from Gene Expression Omnibus and processed. All 232 .`cel` files under GEO accessions GSE33790, GSE33996, GSE5620, GSE5621, GSE5622, GSE5623, GSE5624, GSE5625, GSE5626, GSE5627 and GSE5628 (see `supplemental_-1_cleaned_cel_file_info.csv` for all details of files used). These files were quantile normalised using RMA (Hobbs et al., 2003) with median polishing. The normalised log-transformed expression data were used as input data for training sets.

### Preparation of training, test and hold-out validation sets from normalised array data

The `AtRegNet` database was downloaded from Agris Knowledgebase, specifically the file - http://agris-knowledgebase.org/Downloads/AtRegNet.zip, from this a list of confirmed, direct regulatory transcription factor and target relationships was extracted - these form the basis of positive training examples. Arabidopsis Genome Initiative codes (AGI) for genes in AtRegNet were mapped to Affyemtrix ATH121501 probesets using information in the TAIR AFFY - AGI mapping in https://www.arabidopsis.org/download_files/Microarrays/Affymetrix/affy_-ATH1_array_elements-2010-12-20.txt. Expression profiles for each pair of TF/target genes were extracted and labelled as positive examples. An equal number of randomly selected pairs of gene expression profiles were selected and labelled as negative training examples. A balanced dataset of 8704 training examples was produced in this way.

### Code and data

All code for preparation of training sets is provided in a code and data repository at https://github.com/danmaclean/tf_cnn.

### Development of deep neural networks

All neural network models were developed using the `keras` library (version 2.2.4) in R (version 3.5.2), an API for the TensorFlow library (version 1.10). All code was developed in RStudio (version 1.1.463) and is provided in the code and data repository accompanying this manuscript.

### Development of web application

The web facing version of the tool was developed using the R `shiny` package (version 1.2.0) and is hosted on `shiny`.io at https://danmaclean.shinyapps.io/query_pairs.

## Results

### Developing a Convolutional Neural Network to classify TF/Target pairs

I extracted expression profiles for the 4351 TF/Target pairs from the normalised expression data, labelled these as positive training examples and generated a further 4351 random pairs of expression profiles, to be labelled as negative training examples. This resulted in a tensor of dimension 8702, 232, 2. The tensor was shuffled, and divide so that 80 % was used for training, 10 % validation and 10 % final hold-out test (6900, 901, 901 profiles per set, respectively).

As the individual data are relatively small (2 × 232 matrices) a small model with few layers was tried. I built a separable CNN with two convolutional layers separated by a Max Pooling layer, the convolutional layers feed into a single dense layer before the final classification layer. The `relu` function was used as the activation in all layers except the final dense which was `sigmoid`. The objective function was `binary crossentropy` and the optimiszer was RMSprop in all runs. Also a batch size of 512 was used. All runs lasted for 30 epochs of training. This basic structure is summarised in Figure 1.

**Figure 1:**
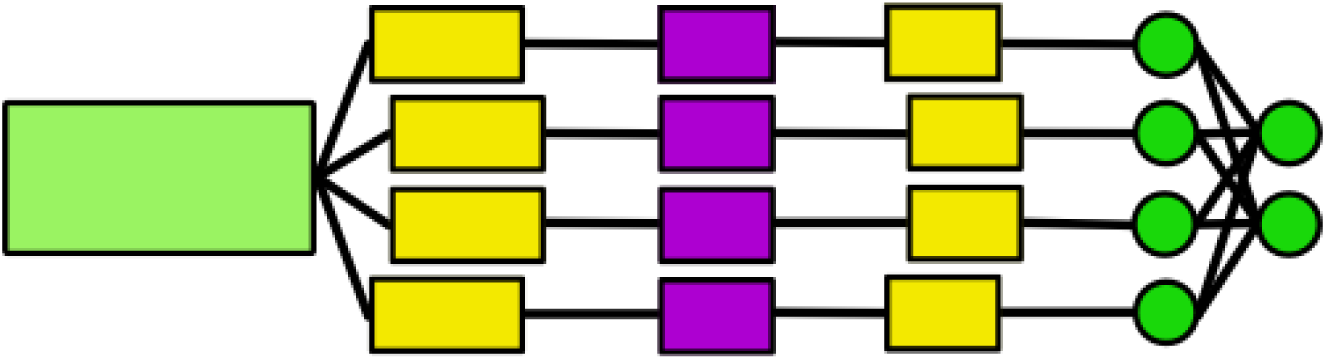
Schematic of initial small CNN model. 2×232 input matrices are fed into a first convolutional layer, a maximum pooling layer, a second convolutional layer then a dense network layer after flattening.

### Estimating hyperparameters

As a first step it was necessary to estimate appropriate hyperparameters of the candidate CNN model. I performed evaluations of the model at different filter and unit counts for the CNN and dense layers. The values were varied through 8,16,32 and 64 and each combination was tested for accuracy on a hold out validation set at the 30th epoch. The accuracy at the 30th epoch is presented in Fig 2. The models each showed accuracy greater than 85% with the highest 91%. In general higher filter and unit counts gave higher hold out validation accuracy, with the highest being the 64/32 filters per convolutional layer and dense layer.

**Figure 2:**
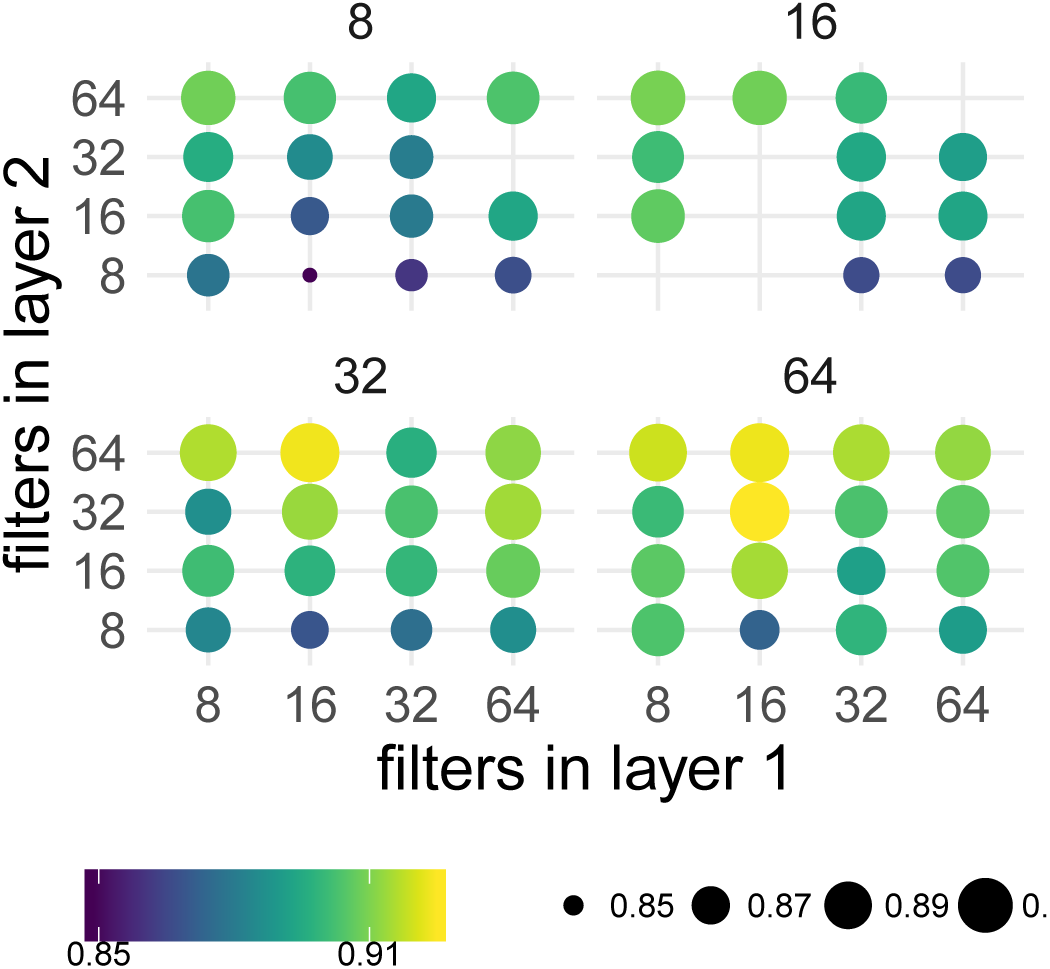
Hyperparameter scan for varied filter and unit counts for a two CNN and one dense layer model. Spot size and colours represent accuracy at the 30th epoch.

On inspection of the training history the individual best models could be seen to be overfitting slightly with increases in loss at the end of the training period (Figure 3 A).

**Figure 3:**
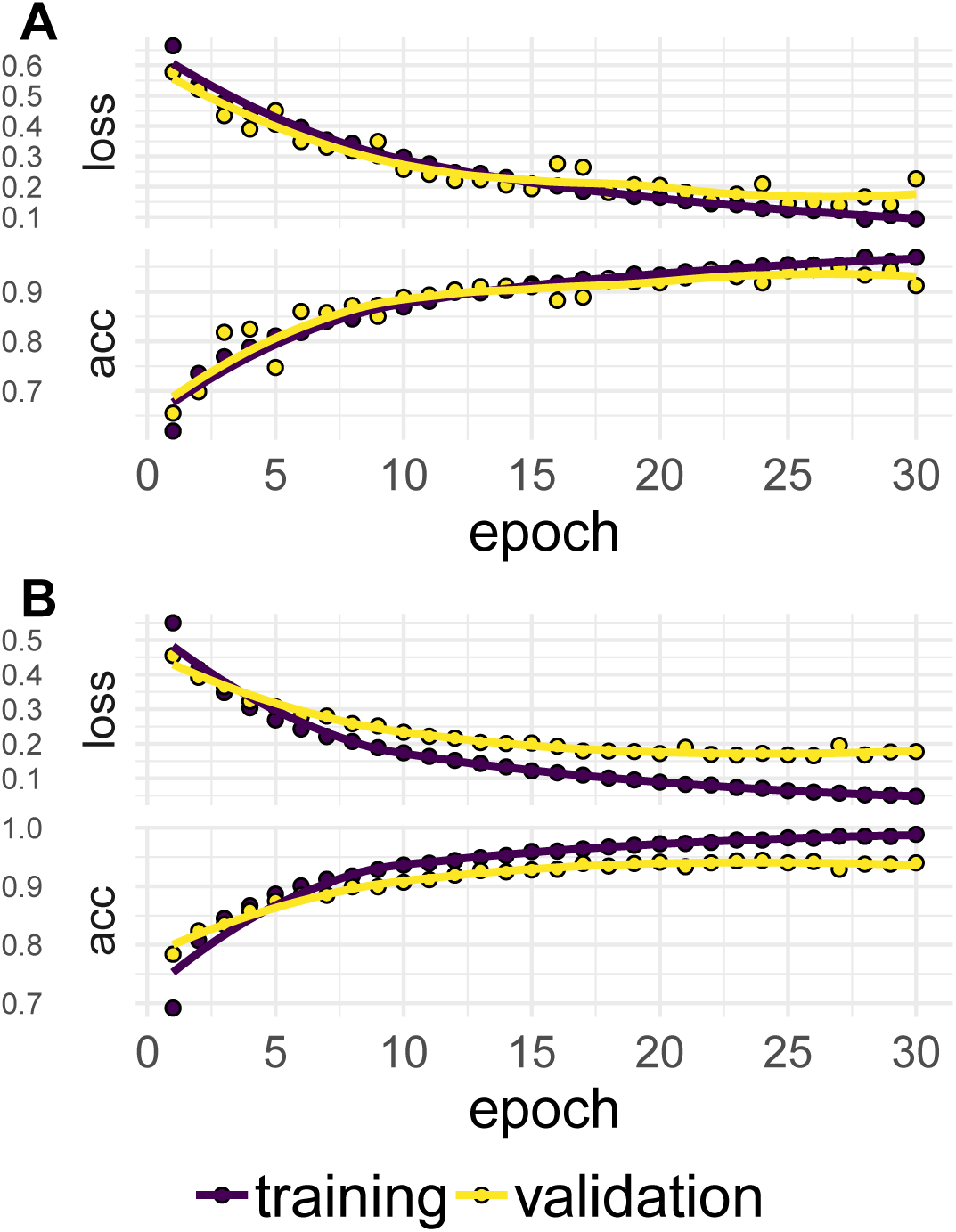
Accuracy and Loss profiles over training history for training and validation set for a model with A) 32,64/32 filters per convolutional layer/dense layer. B) 4,8/4 in the respective layers.

Hence, I selected and tested smaller models manually with slightly lower accuracies to move into a next tuning phase to investigate the effect of regularizations on the accuracy. I made a new base model of 4,8 filters in the convolutional layers and 4 in the dense layer as this showed an accuracy of 0.89 without indication of overtraining.

To finally tune this model I ran further iterations run adding or leaving out batch normalisation layers after convolutional layers and dropout layers after dense layers. I also varied batch size and epoch number. Table 1 show the result and configuration for the top 5 most accurate runs.

**Table 1:** Top 5 runs of a small CNN network model with optional added batch normalisation and dropout layers showing configurations and values.

| Val Acc | Val Loss | Norm Layer 1 Used | Norm Layer 2 Used | Dropout Used | Dropout Rate | Batch Size | Epochs |
| --- | --- | --- | --- | --- | --- | --- | --- |
| 0.9568 | 0.1311 | FALSE | TRUE | TRUE | 0.01 | 512 | 30 |
| 0.9545 | 0.1282 | FALSE | TRUE | FALSE | 0.01 | 512 | 40 |
| 0.9534 | 0.1147 | TRUE | TRUE | TRUE | 0.05 | 512 | 30 |
| 0.9512 | 0.1424 | FALSE | TRUE | TRUE | 0.01 | 256 | 30 |
| 0.9512 | 0.1464 | FALSE | TRUE | TRUE | 0.10 | 256 | 40 |

The most accurate run showed no indication of overtraining at 30 epochs (Figure 3 B. I selected this as a final model and evaluated on the heretofore unseen hold-out validation set for which it showed accuracy of 0.95 and loss of 0.137.

### Distribution of classification probabilities across the entire training set

A useful model would be one that had a clear separation of classifications for positive and negative examples, to understand this for my model I examined the distributions of classification probabilities. I supplied the model with the training data and used it to return probabilities that each training example was a positive example and cross referenced this with the actual class. In the vast majority of cases the classification probability was very close to 0 for negative training examples: 82 percent of points were less than 0.05. Similarly classification probability was very close to 1 for positive training examples: 86 percent of points were greater than 0.95 (Figure 4).

**Figure 4:**
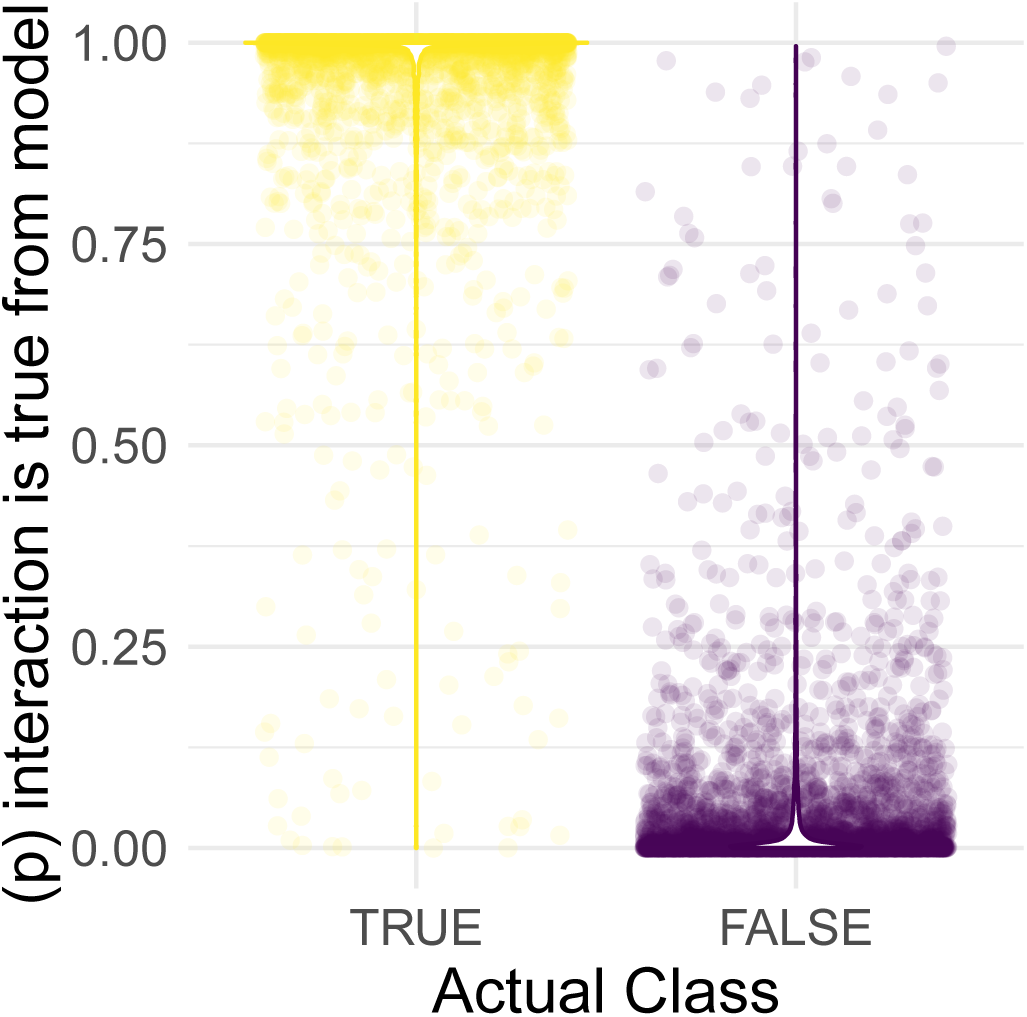
Distribution of model generated probabilities of training examples being a true interacting pair

### Assessing the signals the model uses to classify

An advantage of CNNs relative to other deep neural networks is their relative interpretability. A CNN can be exploited to extract activation maps when provided with data, these maps can highlight the regions of the input to which the CNN is most strongly responding - IE which portion of the expression profiles the model is classifying with. To examine the responses of the model, I ran the training data back through and extracted network activations. These profiles were smoothed and the single largest peak per profile extracted. The expression estimates corresponding to these peaks were extracted and clustered. Numbers of clusters were estimated using principal components analysis and clustering performed with *k*-means clustering, *k* = 3, Figure 5 A. The mean cluster profiles were extracted and can be seen in Figure 5 B. All the mean profiles show a positive correlation between the target and TF, indication that this model is classifying based on positively correlated changes in transcript abundance between target and TF.

**Figure 5:**
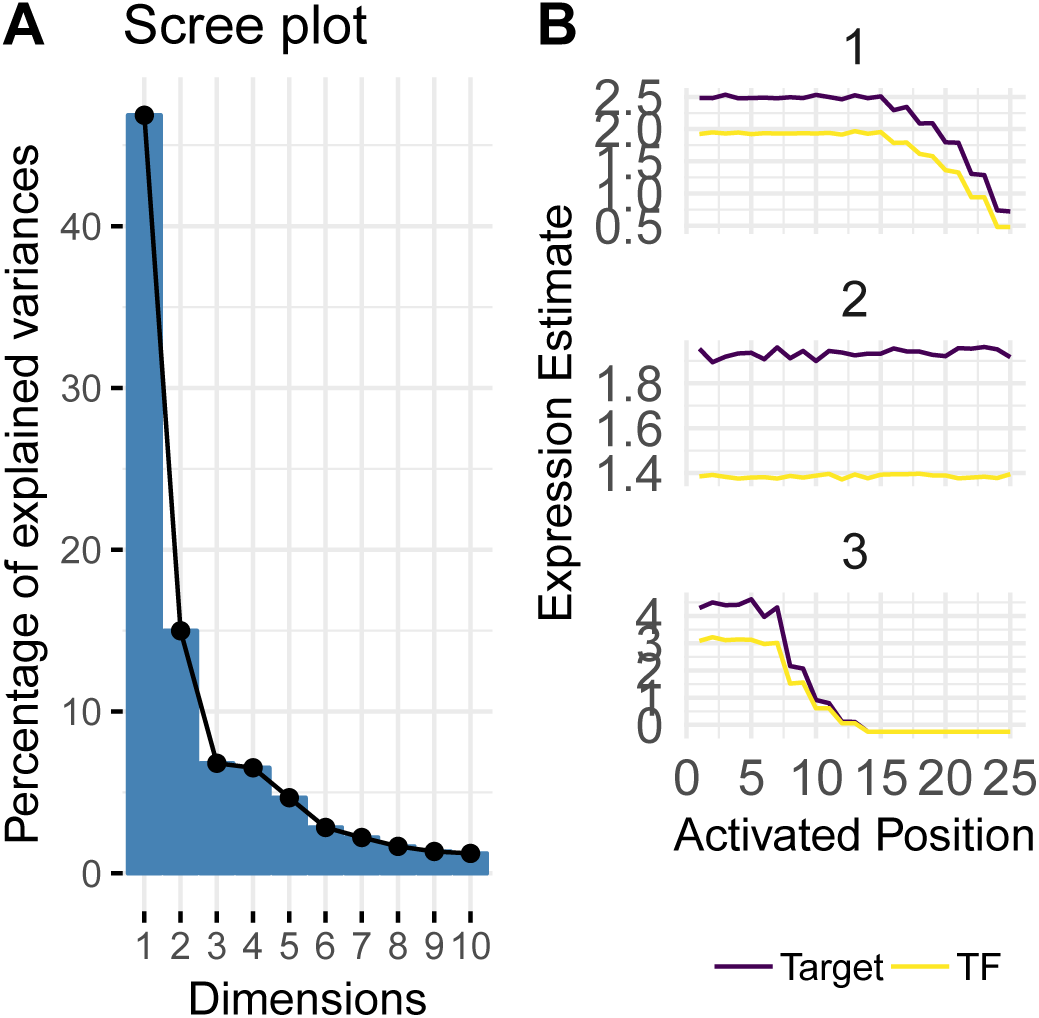
A) Scree plot following principal component analysis of clustered TF/Gene expression profile in regions of highest CNN activation. B) Mean expression profiles of k-means clusters with expression profile extracted from region of highest CNN activation

### A webtool incorporating the model so that can be easily used for prediction

To make the tool directly useful to researchers without need for developing code, I created a small webtool that takes gene lists as input and returns the probability of regulation according to the model. This model is available at https://danmaclean.shinyapps.io/query_pairs. The use can supply a list of AGI or Affy format identifiers as potential regulators and as potential target genes. The tool will then apply the model and return a table indicating the prediction probability of each interaction. A figure showing the distribution of the interaction probabilities relative to the training set used in this study is also generated. The user can download the table in spreadsheet friendly format.

## Discussion

In this study I trained and tuned a convolutional neural network using Affymetrix microarray data from *Arabidopsis* plants subjected to abiotic stress. The network was trained on expression profile data in a series of small 2 × 232 matrices. The network trained quickly, in under three minutes on a common laptop configuration. With tuning and hyperparameter optimisation the network achieved accuracy of 95% on around 6000 training examples without overfitting. The produced model seemed to be classifying on positive correlations within the expression profiles. This makes the model a useful tool, in a certain niche. Those who are interested in predicting positive *Arabidopsis* TF and target relationships in abiotic stress would find it useful.

The model is limited and some caveats should be taken seriously. The primary thing to note is that the activiation patterns I observed were positive, and the model appears only to classify on these. Negatively correlated patterns will not be predicted as true classes. A true negative regulator of expression will probably not classify with the model.

A behaviour like this is probably inherited from the training set. Positively correlated interactions would seem to be the ones most easy to discover and verify and therefore the most numerous sort of interaction in the biological experiments from which the interaction data were distilled.

The model could be extended, the relatively small AtGenExpress expression dataset could be replaced by larger more sensitive and expansive RNAseq datasets for input expression profiles.

The most important aspect of the work presented here is that it was straightforward to build and optimise an effective deep learning classifier on a small-ish training set (thousands rather than millions of training examples) without large compute resources. The work is an excellent example of how datasets generated by individual laboratories could be utilised in deep learning. By taking the optimisation strategy and model structure described here as a starting point many small local datasets could be put to use in myriad classification exercises.

